# Aggressive or moderate drug therapy for infectious diseases? Trade-offs between different treatment goals at the individual and population levels

**DOI:** 10.1101/307223

**Authors:** Jérémie Scire, Nathanaël Hozé, Hildegard Uecker

## Abstract

Antimicrobial resistance is one of the major public health threats of the 21^st^ century. There is a pressing need to adopt more efficient treatment strategies in order to prevent the emergence and spread of resistant strains. The common approach is to treat patients with high drug doses, both to clear the infection quickly and to reduce the risk of *de novo* resistance. Recently, several studies have argued that, at least in some cases, low-dose treatments could be more suitable to reduce the within-host emergence of antimicrobial resistance. However, the choice of a drug dose may have consequences at the population level, which has received little attention so far.

Here, we study the influence of the drug dose on resistance and disease management at the host and population levels. We develop a nested two-strain model and unravel trade-offs in treatment benefits between an individual and the community. We use several measures to evaluate the benefits of any dose choice. Two measures focus on the emergence of resistance, at the host level and at the population level. The other two focus on the overall treatment success: the outbreak probability and the disease burden. We find that different measures can suggest different dosing strategies. In particular, we identify situations where low doses minimize the risk of emergence of resistance at the individual level, while high or intermediate doses prove most beneficial to improve the treatment efficiency or even to reduce the risk of resistance in the population.

**Author summary:** The obvious goals of antimicrobial drug therapy are rapid patient recovery and low disease prevalence in the population. However, achieving these goals is complicated by the rapid evolution and spread of antimicrobial resistance. A sustainable treatment strategy needs to account for the risk of resistance and keep it in check. One parameter of treatment is the drug dosage, which can vary within certain limits. It has been proposed that lower doses may, in some cases, be more suitable than higher doses to reduce the risk of resistance evolution in any one patient. However, if lower doses prolong the period of infectiousness, such a strategy has consequences for the pathogen dynamics of both strains at the population level. Here, we set up a nested model of within-host and between-host dynamics for an acute self-limiting infection. We explore the consequences of drug dosing on several measures of treatment success: the risk of resistance at the individual and population levels and the outbreak probability and the disease burden of an epidemic. Our analysis shows that trade-offs may exist between optimal treatments under these various criteria. The criterion given most weight in the decision process ultimately depends on the disease and population under consideration.

## Introduction

Drug resistance is a rising source of concern for health organizations worldwide [1,2]. The introduction of new antimicrobial agents is often followed closely or even preceded by the emergence of resistant strains, which threatens the efficiency of treatments [3,4]. It is therefore necessary to adopt treatment strategies that successfully treat diseases while minimizing the risk of resistance.

One component of treatments that can be varied is the drug dose (for a review, see [5]). A lower bound is given by the minimal dose that provides a sufficient therapeutic effect. The upper limit for the dose is set to prevent toxic drug effects. Together, these two boundaries define the therapeutic window. For decades, the commonly accepted strategy has been to treat infections as harshly as possible, that is to choose the highest non-toxic dose [6,7]. On the one hand, this choice is driven by the expectation that a high dose leads to a rapid eradication of the pathogen, enhancing the chances of survival and swift recovery of the patient. With respect to the evolution of resistance, a high dose could prove beneficial for two reasons. First, a rapid elimination of the sensitive strain reduces the number of new resistance mutations occurring during treatment. Second, for a sufficiently high dose - above the mutant prevention concentration [8] - resistance might be several mutational steps away, making the emergence of resistance considerably less likely. Also, if the dose is so high that resistance is physiologically impossible, no resistance problem exists.

The probability that resistance evolves does not only depend on the rate at which mutations appear but also on the establishment probability of the resistant strain. A high drug dose limits the rate of appearance. However, it might facilitate the proliferation of resistant mutants. The concern is that the elimination of the sensitive strain allows the resistant strain to spread more easily by releasing it from the pressure of competition. This is particularly risky if resistant types pre-exist prior to treatment. Moreover, the immune response might not be strongly triggered if the drug-sensitive pathogens get quickly cleared by the drug. Resistant pathogens are therefore not killed by the immune system while rare, potentially allowing them to reproduce and reach high numbers. These considerations have led several authors to question the standard “hit hard” strategy and to suggest that – at least under some circumstances – a milder treatment might be preferable [9–13]. Experimentally, the idea of low-dose treatment has received particular attention for the treatment of malaria infections [14–17]. For example, Huijben et al. (2013) find that for rodent malaria, low drug doses reduce resistance with no cost in terms of host health [14].

Combining all factors, the adopted schematic view is that the probability of resistance emergence follows an inverted U-curve [5,18–26] (but more complicated shapes are conceivable): for very low doses, there is no selection for resistance, and for very high doses, resistance is either unlikely due to a large genetic barrier or even physiologically impossible. Kouyos et al. (2014) and Day and Read (2016) argue that, eventually, the range of the therapeutic window determines whether a harsh or a mild treatment is best to mitigate the evolution of resistance [5, 12].

An aspect that has received surprisingly little attention so far is the effect of dosage on the disease dynamics in a community [5]. Virtually all studies on drug dosage focus on the evolution of resistance within a single patient (e.g. [10,12,14,27–29]; for an exception see [30]). However, for infectious diseases, it is equally (or even more) important to constrain the transmission of resistance, and it is not clear whether a dose best at reducing the probability of resistance within a single host is optimal for managing the disease in the population.

Population level considerations resemble those at the within-host level. Again, two factors are crucial for the evolution of resistance: the total rate at which patients develop a resistant infection, and the probability that an existing resistant strain spreads across the community. With a mild treatment, the infection duration may be prolonged, increasing the risk to infect another member of the community and hence ultimately the total number of infections. Each of these infections might, in turn, become resistant. It is therefore conceivable that even if a mild treatment reduces the risk of *de novo* resistance in every single patient, it increases the rate of resistance appearance in the population. On the other hand, a high dose treatment could lead to a competitive release effect at the between-host level, allowing the resistant strain to spread more easily. Colijn and Cohen (2015) study the effect of dosage on the level of resistance in a between-host model [30]. Performing a parameter sensitivity analysis, they show that competition between strains is indeed crucial for whether a mild or a harsh treatment reduces levels of resistance. However, Colijn and Cohen (2015) do not couple the epidemiological model with any within-host dynamics. It is hence not possible to see whether trade-offs between the two levels exist.

Also, managing resistance is not the only treatment goal at the population level (nor is it at the level of the individual patient). At the same time, the spread of the sensitive strain needs to be constrained.

In this article, we set up a nested model, taking into account both the dynamics within patients and the spread of the disease between individuals. Resistance mutations can appear *de novo* within patients and can also be transmitted between individuals. For a more intuitive understanding of the results, we complementarily analyse a standard epidemiological model, for which we estimate the parameters from stochastic within-host simulations. To determine optimal dosing, we apply different measures of treatment success: (1) emergence of resistance at the individual and (2) at the population level, (3) the outbreak probability of an epidemic, and (4) the total disease burden in the population. We find that, in many cases, the choice of a treatment strategy entails trade-offs between these different criteria. For instance, low dose treatment might minimize the probability of emergence of resistance at the within-host level but maximize transmission of resistance at the between-host level. This demonstrates the importance of considering both scales and several criteria for determining optimal dosage.

## Result

### Conceptual framewor

Treatment of mild self-limiting infections is considered by some to be one of the main drivers of the spread of resistance in the community [31]. For this reason, we model an acute self-limiting disease, such as acute otitis media, strep throat or tonsillitis, where the immune system is able to clear the infection even in the absence of the drug [13,31]. Drug treatment speeds up recovery and reduces the infectious period. The effects of treatment depend on the dose *c*. Resistant pathogens can withstand considerably higher drug doses than sensitive pathogens. Treatment is administered once symptoms set in. Symptoms are caused directly by the pathogen and appear when the pathogen load reaches a given threshold. The strength of the immune response is measured in our model by a single variable *I* that is increased proportionally to the number of pathogens inside the host. The immune system does not discriminate between the two strains. It is the only force besides treatment that limits the growth of the pathogen population (no resource competition).

Initially, a single individual is infected with the sensitive strain. How well the infection spreads is determined by the basic reproductive number, defined as the mean number of secondary cases caused by one infectious host in an otherwise fully susceptible population. This number crucially depends on the length of the infectious period and thereby on the drug dose. In the following, we denote by *R*_0_ the basic reproductive number of the sensitive strain in the absence of treatment. In the model, infection happens randomly between hosts with some transmission coefficient *β* (*R*_0_ is proportional to *β*). New infections occur at rate *βSI*, where *S* and *I* denote the number of susceptible and infectious hosts, respectively. We assume that a single strain type is transmitted at each infection event. This is based on the idea that the inoculum consists of pathogens from the same spatial location and that pathogens in close spatial proximity derive from the same clone. Recovered individuals are immune to new infection. As a consequence, the epidemic eventually comes to a halt due to a lack of susceptible hosts in sufficient numbers.

To analyse the model, we perform stochastic agent-based simulations of the precise nested within-host and between-host dynamics. We additionally analyse a deterministic epidemic model with susceptible, infected, and recovered hosts (SIR model), for which we estimate the parameters from stochastic within-host simulations. Details on the model, the analysis, and the precise definitions of the measures used to evaluate treatment success are given in the Methods section.

Fig 1 conceptually summarises the various dose-dependent forces and factors that affect the dynamics of the resistant strain at the individual and population levels.

**Fig 1.**
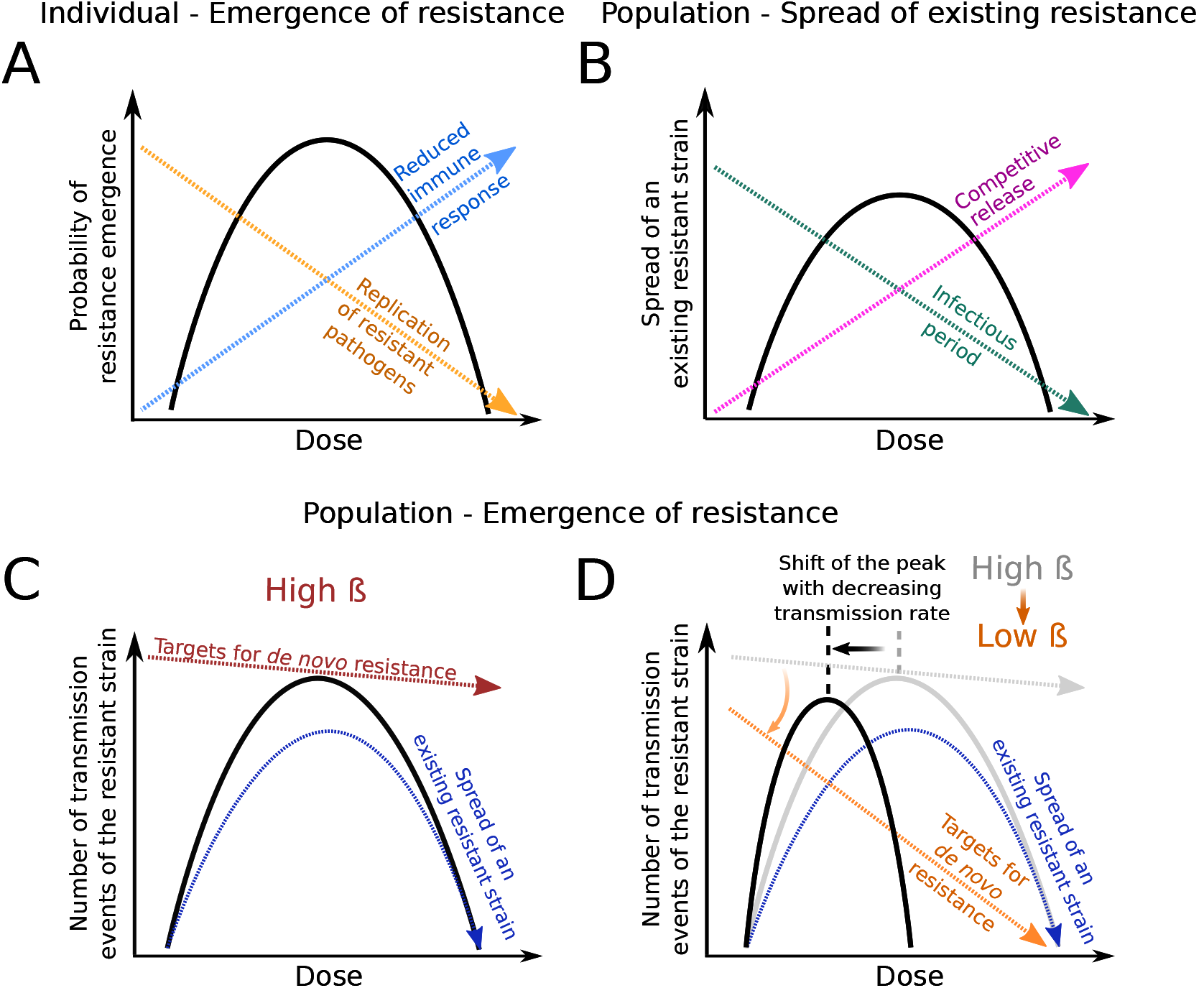
Risk of resistance and the factors shaping it at the individual and population levels. (**A**) Within-host probability of resistance emergence. With increasing drug pressure, the immune response triggered by the sensitive strain is weaker (dashed blue arrow) and resistant pathogens are therefore more likely to establish, but this advantage is counteracted by an increase in pathogen clearance (even of the resistant strain) through the drug (dashed orange arrow). The within-host probability of resistance emergence as a function of dose is shaped by these two opposing forces, and as a result, has an inverted “U-shape” (solid black line). Panel adapted from Kouyos et al. [5]. (**B**) Spread of an existing resistant strain in a susceptible population. Here, increased drug pressure favors the spread of a resistant strain through competitive release (dashed pink arrow), since more susceptibles remain untouched by the sensitive strain. However, with increasing dose, the infectious period (dashed green arrow) decreases, even for infections with the resistant strain. The resistant strain therefore causes fewer secondary infections. Consequently, the curve that represents the spread of an existing resistant strain as a function of the drug dose (solid black line) also has an inverted U-shape. (**C,D**) Number of transmission events of the resistant strain, at a high (C) or low (D) transmission rate. This measure (solid black lines) depends on the ability of a resistant strain to spread (dashed dark-blue arrows, see panel B), and on the rate of *de novo* development of a resistant infection. This latter rate depends on the number of sensitively infected patients (or “targets for *de novo* resistance”), which in turn depends not only on the drug dose but also on the transmission coefficient. In a high transmission setting, large outbreaks of the sensitive strain guarantee a high probability of emergence of resistance (red dashed line in Fig 1C). In contrast, in a low transmission setting, this probability severely decreases with increasing drug dosage (dashed orange arrow in Fig 1D). The location of the peak in the number of transmission events of the resistance therefore depends on the transmission coefficient of the disease. The black curve is essentially the product of the red (or orange) arrow with the dark-blue arrow. The grey line and arrow in panel D are a repetition of the red arrow and black line in panel C.

At the within-host level, the factors that shape the probability of resistance emergence are 1) the direct effect of the drug on the resistant strain and 2) the suppression of the resistant strain through the host’s immune response (Fig 1A). At the population level, the factors that determine how well the resistant strain spreads once it has emerged within a host are the infectious period for infections with this strain (directly affecting the number of secondary cases) and the strength of competition for susceptible hosts with the sensitive strain (Fig 1B). As explained in the detailed model analysis, in both cases, these antagonistic dose effects lead to an inverted U-curve, and we argue below that the maximum in each of these curves lies at the same dose.

However, in order to assess the risk of resistance in the population, it is not sufficient to consider the spread of resistance once it has emerged. Another key factor is the rate at which resistance emerges *de novo* in the population. This rate does not only depend on the within-host probability of resistance but also on the number of sensitively infected hosts who might develop a resistant infection. This number in turn depends on the drug dose but also on the transmission coefficient of the disease in the population. Figs 1C and D anticipate a primary insight: the number of transmission events of the resistant strain (as a measure for the risk of resistance in the population) displays a single maximum, the location of which depends on the transmission coefficient of the disease. Thus, the curves describing the risk of resistance at the individual and population levels may peak at very different drug doses, meaning that the optimal dosage to combat resistance in a population cannot be directly inferred from the optimal dosage to combat resistance inside a single host.

In the following, we provide a quantitative model analysis to support the statements made in the previous paragraphs. Thereafter we also explore the effect of drug dose on two other population-level measures of treatment success (outbreak probability and disease burden). We put the focus on uncovering trade-offs between optimizing treatment benefits under one or the other measure.

### Within-host probability of resistance emergenc

Our within-host model of an acute, self-limiting infection was adopted from the theoretical study by Day and Read [12]. They showed that the probability that resistance evolves *de novo* within a single patient, *p_e_*(*c*), follows an inverted-U-shaped curve with a single maximum (see Fig 1A and 2A). For low doses, the sensitive strain triggers a strong immune response, and resistant pathogens are likely to be suppressed by the immune system before reaching high levels. For high doses, resistance is not sufficient to withstand the drug. The maximum corresponds to doses for which the sensitive strain is cleared too rapidly by the drug to evoke a strong immune response, while the resistant strain is not substantially affected by the drug. (The higher number of *de novo* mutations during treatment with lower doses does not seem to have a major effect on the shape of *p_e_*(*c*) for the chosen parameter set).

**Fig 2.**
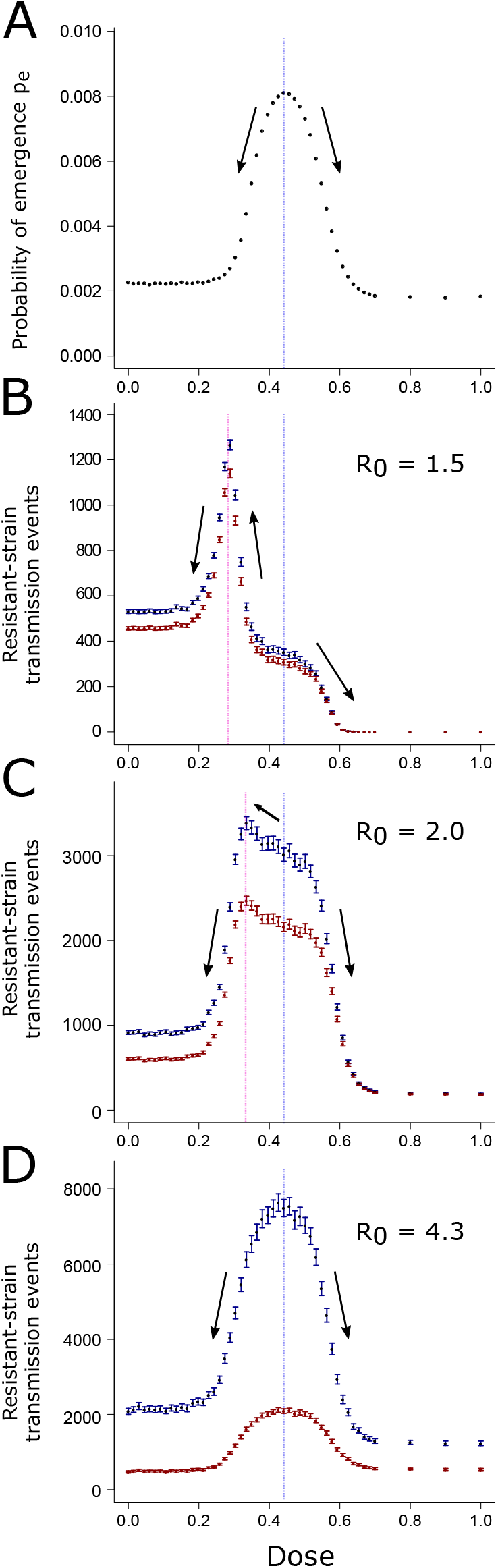
Comparison of optimal dosage determined through the within-host probability of resistance (A) or through the number of transmission events of the resistant strain during an epidemic (B-D). Blue symbols show the total number of transmission events, while red symbols show only transmission events towards susceptible hosts. The vertical blue line corresponds to the peak in the within-host probability of resistance and the pink line to the peak in the number of resistant-strain transmission events. The black arrows represent the trend of the measure of interest when one shifts the treatment dosage away from the dose giving a maximum in *p_e_*. Mean over 2, 800 to 40,000 simulations runs per data point. In dark blue and dark red: the 95% confidence intervals of the mean for each dose.

The best dose choice to reduce the within-host probability of emergence is not straightforward. Depending on the therapeutic window (i.e. on the range of doses that can practically be administered), the best dose choice can either be on the left-hand side of the central peak, in the low-dose region; or it can be on the right-hand side in the high-dose one. In any case, as highlighted by Day and Read [12], the optimal dose with respect to this criterion can be nothing else than one of the two boundaries of the therapeutic window.

### Transmission of the resistant strain in the population

We next explore how the expected number of transmission events of the resistant strain in the population, 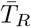, depends on the drug dose (Fig 1C and 2B-D). For all values of *R*_0_,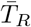is highest for intermediate doses but the location of the peak depends on the transmission coefficient.

In particular, with a rather low *R*_0_ (Fig 2B), the peak in the number of transmission events of the resistant strain is considerably shifted towards lower doses compared to the peak in the probability of within-host emergence of resistance, *p_e_* (shown in Fig 2A). Therefore, there is a non-negligible range of doses for which lowering the dose in order to reduce the probability of within-host emergence of resistance will in return substantially increase the average number of transmission events of the resistant strain. For many if not all strategies for resistance mitigation, this would be the exact opposite of the aimed outcome. For increasing transmission coefficients (Fig 2C,D), the peak in the number of transmission events of the resistant strain converges to similar doses as the peak in *p_e_*. This means that, for high *R*_0_,decisions made to minimize *p_e_* will also minimize the number of events of transmission of the resistant strain and vice versa. If we consider transmission events towards susceptibles only (red symbols in Fig 2), the same conclusions for the location of the peak hold. However, the peak is then highest for intermediate *R*_0_, presumably because for high *R*_0_, too few targets of infection are available.

What explains the difference in the location of the peak for low and high drug doses? The number of transmission events of the resistant strain, *T_R_*, is influenced by two factors: the rate at which the resistant strain appears in the population (i.e. by the rate at which some host develops a resistant infection) and its chances to spread once it has appeared.

These two factors need not peak simultaneously. The rate at which the resistant strain appears in the population is determined by how likely the resistant strain is to emerge within a host (approximately given by *p_e_*) and by the number of hosts infected with the wild-type strain. The more individuals are infected by the wild-type pathogen, the more chances the resistant strain has to emerge. On the other hand, large wild-type outbreaks that favor the *de novo* appearance of the resistant strain in the population do not provide ideal conditions for its spread. For large wild-type outbreaks, by the time the resistant strain appears, many individuals have already acquired immunity through infection with the sensitive strain. At low doses, where the sensitive strain is able to infect many individuals, the spread of the resistant strain is hence hindered through the greatly reduced pool of targets of infection and through competition with the sensitive strain for the remaining susceptibles. At high doses, it cannot spread well either since it has an equally low fitness as the sensitive strain. Spread of the resistant strain is hence expected to be easiest at intermediate drug doses, where it experiences competitive release from the sensitive strain and is itself barely affected by treatment. Indeed, it seems intuitive that spread is easiest at similar doses as the spread of the resistant strain at the within-host level (which largely determines the peak in *p_e_*): the differential recovery rates for patients who are infected with the sensitive or the resistant strain reflect the differential growth rates at the within-host level (compare Fig 8A and B); the former strongly affect the spread of the resistant strain at the population level, the latter at the within-host level.

For a high *R*_0_, a large initial epidemic of the wild-type pathogen is likely even for high drug doses. A large wild-type epidemic ensures that the resistant strain appears in some hosts in the population. Therefore, for high *R*_0_, the shape of 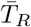is dominated by the rate of spread (Fig 2D). In contrast, for low *R*_0_, appearance of resistance at the population level is a limiting factor.

The probability of a large outbreak (with a large opportunity for resistance to appear) drops strongly with the drug dose, shifting the peak to lower doses, where large outbreaks are more likely (Fig 2B; cf. also Fig 1C). Note that the shift in the peak is primarily due to stochasticity in the appearance of resistance. This includes in particular that resistance might not appear at all before extinction of the disease. In the classical epidemic model with susceptible, infected, and recovered hosts (SIR model, see Methods section), the integral 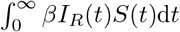quantifies transmission of the resistant strain (there is no superinfection in the SIR model and all transmission is towards susceptibles). Since our SIR model implementation is deterministic, appearance of the resistant strain is guaranteed. Then, the location of the peak depends only weakly on *R*_0_(not shown).

In section S5, we analyze an endemic rather than an epidemic disease, modeled by an SIR model with immigration/birth and death. We assume that the population is at endemic equilibrium prior to the evolution of resistance and evaluate the risk of resistance emergence. The model allows for an analytical treatment, making it easy to disentangle the two factors contributing to resistance (rate of appearance and spread). We find similar patterns in the risk of resistance as for the epidemic disease considered in the main text. In particular, the risk of resistance in the population can peak for considerably lower doses than the within-host emergence of resistance (Fig S7). This also holds if immunity is not life-long but lost at a certain rate (Fig S8).

### Outbreak probabilit

Initially, only a single individual is infected. It is therefore possible that by chance, the disease disappears from the population after just a few infection events. If it survives this initial phase, the population is hit by an epidemic outbreak. The stronger the dose administered to the infected patients in the population, the less likely an outbreak is to occur (Fig 3). The first few infected individuals are very likely to be mostly carrying the sensitive strain. A stronger dose decreases the time during which these individuals are infectious, making them less likely to transmit the pathogen further in the population (cf. also SI section S2). This in turn reduces the outbreak probability. Thus, the increased recovery rate in early-infected patients associated with an increased treatment strength is key in reducing the likelihood that a budding epidemic can develop further. Therefore, to reduce the outbreak probability, one should prescribe the highest available dose of treatment to early-infected patients. As can be seen in Fig 3, the relative reduction in the outbreak probability for high compared to low doses is stronger for low than for high *R*_0_.

**Fig 3.**
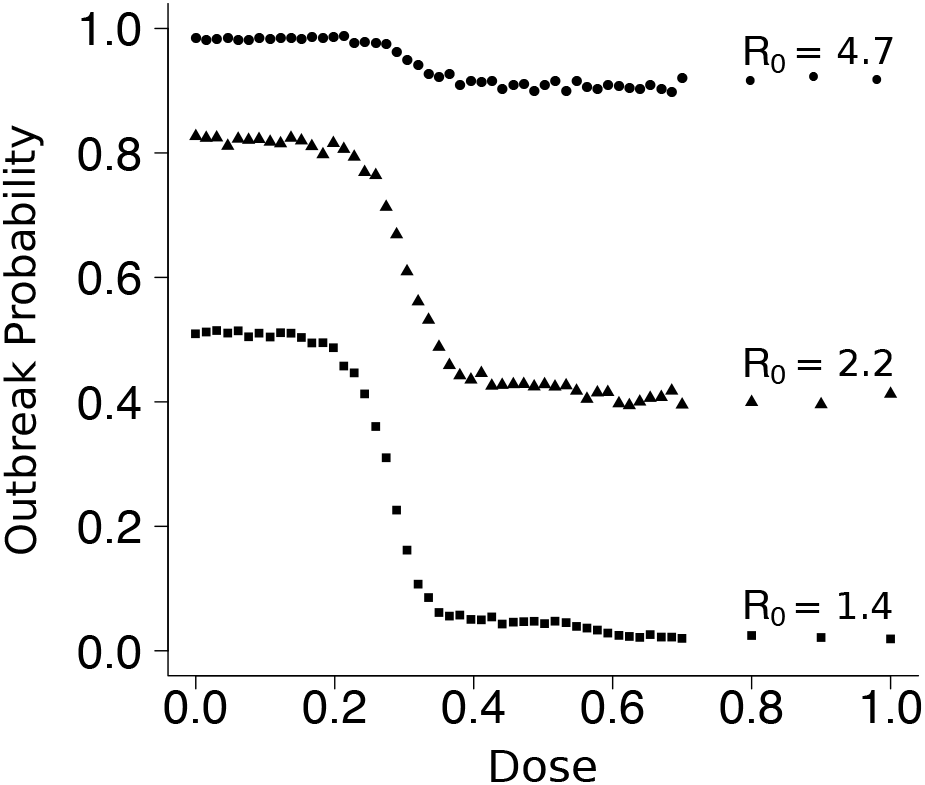
Effect of the treatment strength on the outbreak probability for various transmission coefficients. Mean over 1900 to 5600 simulations per dose and transmission coefficient. Diamonds: *R*_0_ = 1.4 (*β* = 1.5 × 10^-5^ days^-1^). Triangles: *R*_0_ = 2.2 (*β* = 2.3 × 10^-5^ days^-1^). Circles: *R*_0_ = 4.7 (*β* = 5.0 × 10^-5^ days^-1^). The confidence intervals are not shown as they are too small to be clearly seen on the plot.

These considerations can be related to the outbreak probability in a one-strain stochastic SIR model, given by 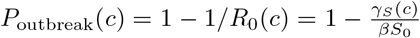 is the dose-dependent recovery rate for infections with the sensitive strain, *β* is the transmission coefficient as before, and *S*_0_ denotes the initial number of susceptible hosts. *P*_outbreak_ shows the same qualitative behavior as observed in Fig 3. However, since the distribution of secondary cases differs between the stochastic SIR model and the agent-based model, the outbreak probabilities differ quantitatively by a large margin (see SI sections S3.4.1 and S3.4.2).

### Disease burden in large outbreak

Last, we consider the disease burden, defined here as an epidemiological measure of the total number of days that individuals spend being infectious (in other words, it can be thought of as (Number of infected people) × (Duration of an infection), see Methods section for details). We focus on the disease burden, provided a large outbreak occurs, and denote this quantity by *B.*

As can be intuitively expected, the overall trend is that the burden *B* decreases for increasing doses (Fig 4). *B* is maximal in the absence of treatment and minimal with very aggressive treatment (where the resistant strain cannot withstand the drug pressure either). Interestingly, however, for intermediate doses, the disease burden displays a local minimum (and maximum) unless *R*_0_ is large (Fig 4A and B). One can already see that, when high drug concentrations are forbidden because they are toxic, the best dose choice to minimize *B* can be the one corresponding to this local minimum. For the following discussion of this result, remember that we only consider outbreaks where the disease burden is at least 300 patient days, meaning that an epidemic outbreak actually occurred. For the majority of cases, this implies a quite large burden of the order of 10^4^ patient days (see Fig S1). Appearance of resistance is hence not limiting, and the population-wide dynamics of the resistant strain are mainly (though not exclusively) determined by its ability to spread. The large number of pathogens of both strains also allows us to draw on the deterministic SIR model, which describes the disease burden surprisingly well (Fig 4A and Fig S5; however, note that the SIR overestimates the depth of the “valley” around the local minimum).

**Fig 4.**
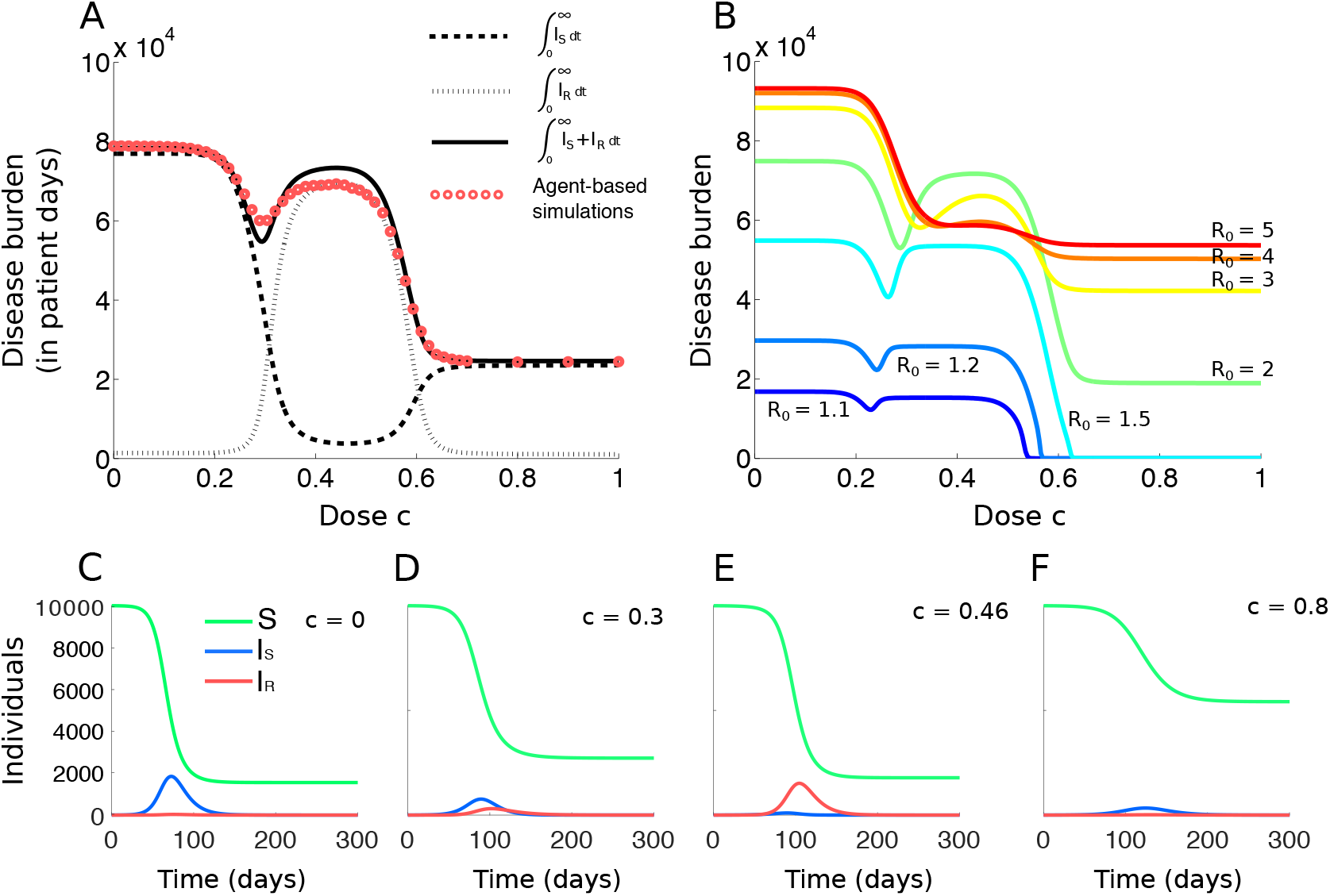
Effect of dose on the disease burden and infection dynamics. A: Comparison of the disease burden obtained with the nested model and the SIR model (*R*_0_ = 2.2). Each red dot represents the mean for 1600 to 4000 runs of simulation of the nested model. For the SIR model, the contributions of the sensitive and resistant strains to the disease burden are also shown. B: Burden curves (in patient days) for various values of Ro. C-F: Exemplary population dynamics of the SIR model for four doses and *R*_0_ = 2.2. Note that the burden *B* is the area-under-the-curve of the sum of the blue and red curves (number of individuals infected by each strain at time *t*).

Fig 4A separates the contributions of the sensitive and the resistant strain to the disease burden, and Fig 4C-F illustrate the disease dynamics of the SIR model for four different drug concentrations. For very low doses, the sensitive strain is barely affected by the drug (Fig 4C). It spreads and as a consequence, the number of susceptible hosts quickly drops below the threshold which would allow the rare resistant strain to spread. The burden caused by the resistant strain is therefore negligible (Fig 4A and C). For slightly larger doses (Fig 4D), as the dose increases, the sensitive strain gets more and more hampered by treatment. However, it is still able to infect a large number of individuals that become immune, and because of the lack of available susceptible individuals, the resistant strain cannot proliferate abundantly in the population. In this region, the disease burden has a minimum (see SI section S7.2 for an analysis with a simplified model). As the drug concentration increases further (Fig 4E), the resistant strain experiences stronger competitive release from the sensitive strain. In this regime, it not only outcompetes the sensitive strain, it also benefits from a large pool of susceptible individuals, left untouched by the sensitive strain due to its low fitness. It hence spreads quickly and causes most of the disease burden in an epidemic nearly as large as in the absence of treatment. For high doses (Fig 4F), where both strains have again similar fitness (i.e. recovery rates), the general dynamics are the same as at very low doses, but neither strain can spread well. Even though the reduction in the burden at the local minimum is not very pronounced, the difference in terms of days spent infected by the individuals of the population can be major in comparison to the neighboring local maxima (in the order of 10^4^ patient days of infection in a population of 10,000).

For large *R*_0_ (Fig 4B), the disease burden as a function of dose is monotonically decreasing. For all doses, the sensitive strain is able to infect too many hosts to allow for a pronounced competitive release effect. Whether the disease burden is monotonically decreasing or displays a minimum at an intermediate dose depends on a complex interplay between the recovery rates and the transmission parameters (see Fig S15 and analytical details in section S7.3).

As a general trend, the burden increases with increasing *R*_0_. A larger transmission coefficient causes, in general, a larger outbreak. However, for intermediate drug concentrations, the disease burden is maximal for intermediate *R*_0_. This means that for such doses, a higher *R*_0_ does not lead to more days of infection as a whole, but to less of them. This occurs for doses around *c* = 0.5 where the competitive release effect is strongest. For large *R*_0_, as discussed above, there is no substantial competitive release, and most individuals get infected by the sensitive strain instead of the resistance strain, contrarily to what happens for these doses at lower *R*_0_. In the intermediate dosing regime, the recovery rate of the sensitive strain is substantially higher than that of the resistant strain. Thus, the disease burden caused by a sensitive-strain epidemic is much lower than the burden caused by a resistant-strain epidemic if intermediate doses are used. This leads to the observed maximum in the burden for intermediate values of *R*_0_ in this dosing regime.

Identical qualitative results are obtained if, instead of considering the disease burden as the number of days members of the population spend being infectious, we restrict the definition of the disease burden to the amount of days individuals spend showing symptoms. In fact, if we focus on symptomatic cases, the “valley” in the disease burden as a function of dose is even deeper than for the infectious burden (see Fig S10). For instance for *β* = 1.9· 10^-5^ days^-1^, the reduction is 25% for the symptomatic burden, while it is only 11% for the infectious burden. The location of the minimum in the burden in dependence of the drug dose remains the same.

Note that we do not observe a non-monotonic behavior in the unconditioned disease burden *B*_0_(c), which considers epidemics of all sizes - including epidemics that die out very early. This is due to the rapid decline in the outbreak probability with increasing dose. (However, we can not entirely exclude that *B*_0_ is non-monotonic as well in certain parameter regions.)

In Fig S11, instead of the disease burden, we consider the total number of individuals infected during the course of the epidemic as a function of the drug dose. This measure also shows a local minimum. For high *R*_0_, this local minimum fades away, similarly as for the disease burden. However, contrarily to the disease burden, the total number of infecteds during the course of an epidemic monotonically increases with *R*_0_ for all doses. This confirms that the larger burden for intermediate *R*_0_ and doses around *c* = 0.5 is attributable to the differential recovery rates of patients infected with the sensitive or the resistant strain. In SI section S7.2, we simplify the true dynamics to two strictly sequential epidemics of the sensitive and the resistant strain and determine the total number of infected individuals of both epidemics. The simplified model reproduces the patterns observed in the full model.

### Trade-offs between treatment goal

From a comparison of Fig 2A and B, we have already seen that a certain dose choice might minimize the risk of *de novo* resistance at the within-host level but maximize the transmission of resistance at the population level and vice versa.

Here, we compare the within-host probability of resistance and the disease burden for three different strategies: (1) using the highest possible dose, (2) using the dose that leads to the lowest within-host probability of emergence *p_e_* (cf. Day and Read, 2016 [12]), and (3) using the dose that leads to the lowest disease burden B. This comparison, shown in Fig 5, is done for three different therapeutic windows. As can be seen, a dosing strategy that is optimal to achieve one goal is suboptimal under another measure of treatment success. E.g., for the first window (Fig 5A), representing a low drug-tolerance scenario, the dose leading to the lowest *p_e_* (orange cross on the figure) leads to the highest disease burden at the population level. Choosing the highest possible dose (pink cross) leads to a suboptimal disease burden as well. For the therapeutic windows in Fig 5B and C, each time, two of the three strategies yield the same outcome, while the third one differs. Other therapeutic windows can lead to even different conclusions, with, for instance, all of these three strategies agreeing on choosing the highest available dose.

**Fig 5.**
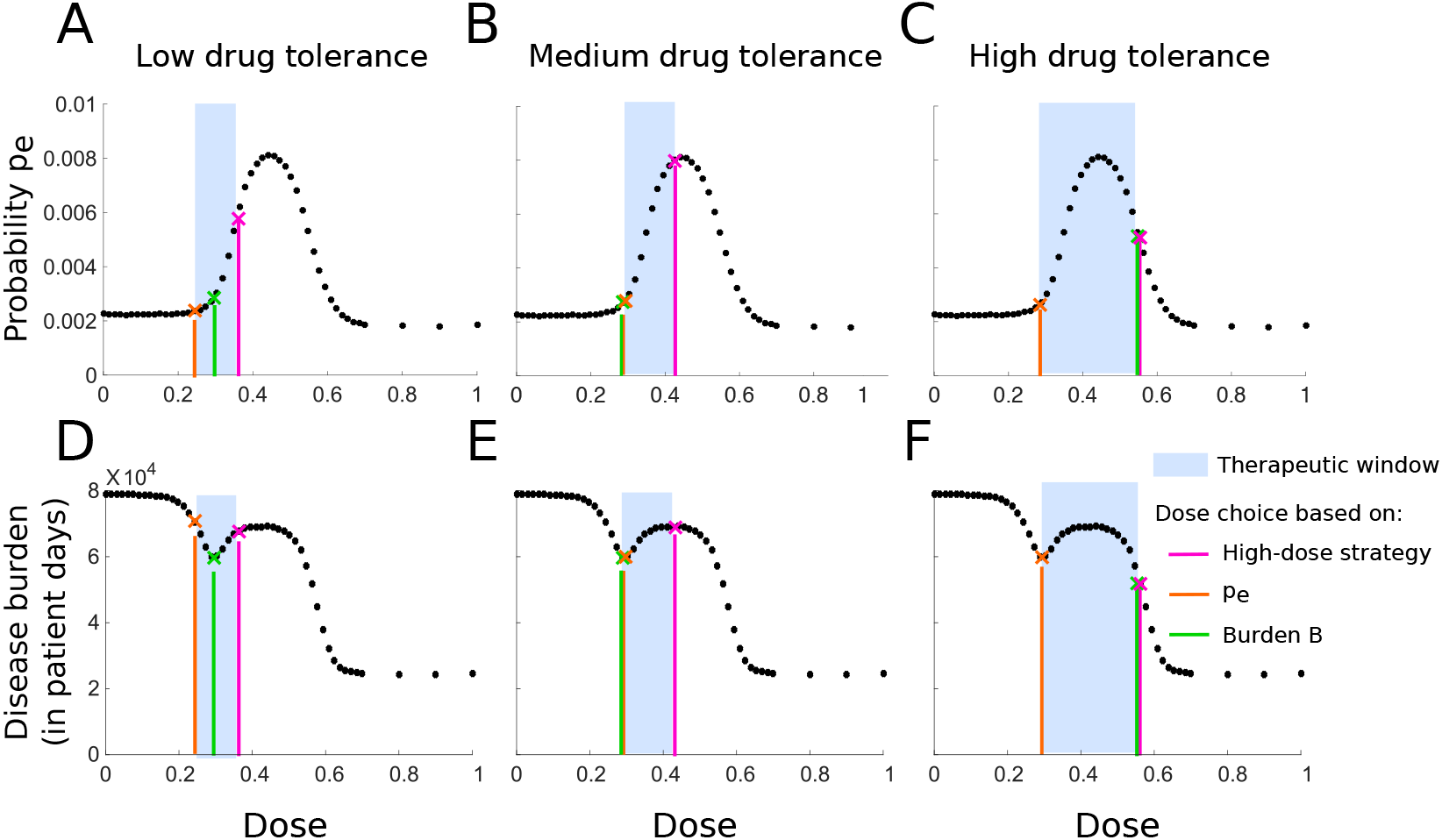
Comparison of three dosing strategies. Strategy that (1) uses the highest possible dose (pink), (2) uses the dose that minimizes the within-host probability of resistance (orange) and (3) uses the dose that minimizes the disease burden (green). Top row: Effect of each strategy on the within-host probability of resistance *p_e_* for three therapeutic windows. Bottom row: Effect of each strategy on the disease burden *B*, where we chose *R*_0_ = 2.2. The three therapeutic windows are: (A, D) low-drug tolerance window (0.25 < *c* < 0.37); (B, E) medium drug-tolerance window (0.32 < *c* < 0.45); (C, F) high drug-tolerance window (0.27 < *c* < 0.55).

### Model extension: Incomplete treatment coverag

The model assumes a perfectly homogeneous population. We now introduce a form of population structure by extending the SIR model to include incomplete treatment coverage. Infected patients receive treatment with probability *f* and remain untreated with probability 1 – *f* (see SI section S4 for the model equations).

For all *R*_0_, the burden as a function of dose becomes monotonically decreasing as coverage gets lower (Fig 6). Therefore, for low treatment coverage, the optimal dose-choice to minimize the burden is the highest possible dose, regardless of where the therapeutic window lies. However, as can be seen in Fig 6A-C for intermediate doses, the disease burden displays an intermediate minimum as a function of treatment coverage. For intermediate doses (roughly 0.3 < *c* < 0.55), treating only a fraction of the population (incomplete treatment coverage) is more beneficial to the population as a whole than treating all infected hosts. In the case of a complete coverage, the resistant strain dominates the pathogen-mix in the population and produces full-fledged epidemics. Conversely, for very low coverage, the sensitive strain dominates. For intermediate coverage, the two strains compete for hosts, resulting in a suboptimal epidemic in total.

**Fig 6.**
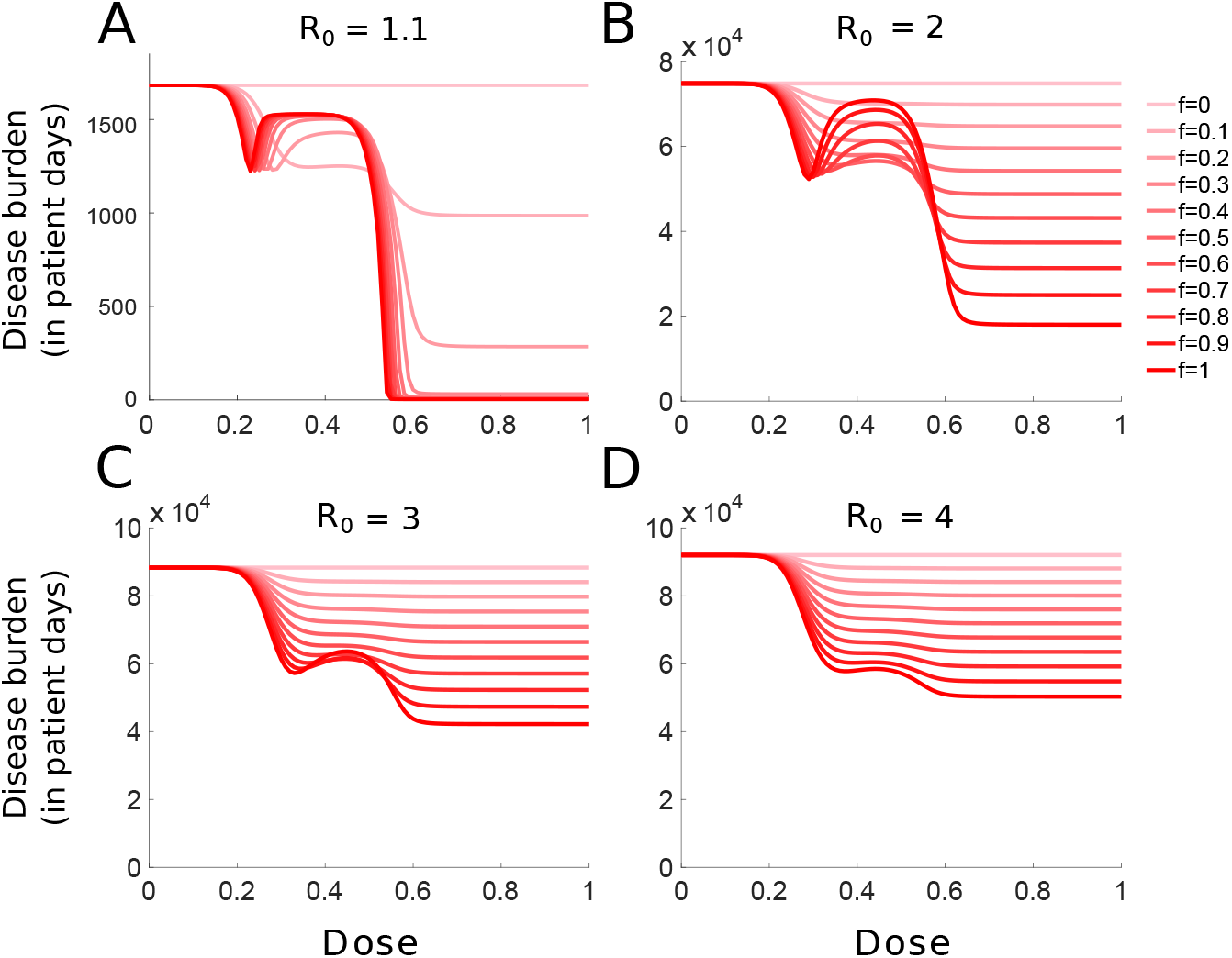
Effect of the treatment coverage on the disease burden. The disease burden for various fractions *f* of treated hosts is shown, ranging from no treatment (*f* = 0, pale red) to full treatment (*f* = 1, dark red). The basic reproductive number is given by: (A) *R*_0_ = 1.1; (B) *R*_0_ = 2; (C) *R*_0_ = 3; (D) *R*_0_ = 4. Note that the scale on the y-axis in panel A is very different from the other panels.

In SI section S5.3, we consider the emergence of resistance in an endemic disease (modeled by an SIR model with birth and death) and find similar non-trivial dependencies on treatment coverage.

## Discussion

From a within-host model, previous work concluded that, in order to minimize the risk of resistance, either the highest tolerable dose or the lowest effective dose should be used [12]. Between these two doses, the right choice depends on where the therapeutic window lies with respect to the “inverted-U-shaped” curve that describes the relationship between the administered dose and the probability of emergence of resistance.

When it comes to infectious diseases, emergence and spread of antimicrobial resistance are not only determined by possibilities for resistance to emerge inside one infected host, but are also strongly influenced by the population-level dynamics of the disease. We therefore used a nested model of within-host and between-host dynamics to study the effect of drug dose on several measures of treatment success at the population level (outbreak probability, transmission of the resistant strain in the population, disease burden).

We showed that these different population-level criteria may lead to different dosage recommendations. What is more, they can strongly differ from recommendations based on measures at the within-host level. We interpreted and expanded our results with the help of simplified epidemiological models. This enabled us to uncover principles that hold true beyond our specific choice of within-host parameters.

### Which dose do the four criteria suggest

The dose choice required to minimize the outbreak probability is the highest non-toxic dose. Therefore, in an ideal case with a perfectly-monitored population, using aggressive antimicrobial chemotherapy on the first few infected patients is best to minimize the probability of an epidemic breaking out.

The optimal strategy to adopt is not as straightforward if the aim is on suppressing antimicrobial resistance. Since the probability of emergence of resistance within one host shows a peak, one must take into account the therapeutic window and choose one of the boundaries to minimize it [12]. But this probability is not enough to assess the consequences of a dose choice at the population level in the case of infectious diseases. The transmission characteristics of the pathogen of interest need be taken into consideration.

Criteria that focus on the emergence and spread of resistance in the population are fundamental for assessing the influence of different treatment strategies on drug resistance. We chose the number of transmission events of the resistant strain as such a criterion. As the within-host probability of resistance, it displays an intermediate maximum. For pathogens with high transmission, the peaks are located at similar doses, and individual-and population-level measures of the risk of resistance lead to similar conclusions. For pathogens with rather low transmission coefficients, in contrast, the peak in the number of resistant transmission events is shifted towards lower doses. In that case, adopting a dose that minimizes the probability of emergence of resistance inside a host may actually maximize the average spread of resistance in the population. Thus, the conclusions reached by measures at the individual and population levels can prove to be dramatically different. These results remain true when relaxing the closed-population assumption of our model by adopting an SIR model with birth and death, in which the population is at the endemic equilibrium prior to the emergence of resistance (see section S5).

As for minimizing the overall disease burden borne by the infected population the best strategy strongly depends on the pathogen’s transmission coefficient. We find that in parameter regimes with low transmission coefficients, a local minimum, maximum, or both, can be observed within the therapeutic window of interest. In that case, the optimal dose to minimize the disease burden can range from the lower bound of the therapeutic window to its higher bound. In particular, intermediary doses inside the therapeutic window can be the most beneficial doses. For higher transmission coefficients, the disease burden decreases monotonically with the treatment strength. Then, aggressive chemotherapy is consistently optimal, as it is optimal for minimizing the outbreak probability.

We also show that incomplete treatment coverage (e.g., due to asymptomatic cases or infecteds not seeking medical help) can alter the optimal dose choice. In particular, the disease burden decreases monotonically for low treatment coverages. For epidemics in which only a small portion of the population can be treated, adopting aggressive chemotherapy can be beneficial to the population as a whole, regardless of the R_ừ_. Moreover, we show that for intermediate doses (which favor the resistant strain), the disease burden displays a local minimum as a function of coverage for low transmission epidemics. There, a partial coverage stimulates competition for hosts between the resistant and sensitive strain, resulting in a smaller epidemic overall (see also [33–36] and discussion further below). It can therefore be beneficial to the population as a whole to actively reduce coverage in cases where the therapeutic window is restricted to these intermediate doses.

Overall, the four criteria do not always agree on what the best dose choice is, and trade-offs exist between different treatment goals, which raises the question of which criterion should be used.

### Which criterion should be used

Obviously, it is most desirable to prevent a large outbreak from occurring in the first place, which is best achieved with high drug doses. However, this requires to detect and to treat all (or at least most) of the early cases, which might not be feasible in reality. Moreover, even with the highest drug dose and perfect treatment of all symptomatic cases, the outbreak probability is still ≈ 40% for the parameter choice of Figure 3. Our intuition (which is supported by the considerations in SI section S2) is that the possibility of resistance generally does not affect the outbreak probability. Resistance only has a small probability to appear in any one host and hence most likely emerges in the population only once a major outbreak has started. However, since people travel (which we do not consider in our model), resistance might be imported from the outside [37]. In this case, using high drug doses is likely to be less efficient in reducing the risk of a large outbreak (see also a corresponding comment in [33]).

When the epidemic cannot be contained and a large outbreak occurs, the criterion that should guide decisions is not obvious and largely dependent on the disease of interest. Preventing the evolution and spread of resistance is not a goal per se (otherwise, not treating at all would be best). Tanaka et al. (2014) distinguish direct and indirect benefits of treatment [38]. Direct effects of treatment refer to the successful therapy of infected individuals, reducing their morbidity (or mortality). Indirect effects focus on the population as a whole and have been quantified based on the total number of cases over the course of the epidemic [33,36–38,39]. The disease burden as a measure combines both of these effects since it is reduced both by the prevention of cases and the successful treatment of infections. The disease burden (restricted to symptomatic cases) can be understood as the total number of days infected individuals of a population spend being incapacitated by the infection. It can therefore be most relevant for diseases that would cause symptoms severe enough to force infected individuals to stop their activities for a few days, but not severe enough to cause any deaths.

The disease burden is a meaningful criterion from an utilitarian point of view. However, it ignores the perspective of any specific patient once treatment is administered. The wish of any patient is to recover as quickly as possible, and physicians usually take their decisions accordingly. We have not included this as a criterion in the main part of the manuscript. However, from a look at the mean recovery rates (Fig 8B) and the low probability of within-host resistance (Fig 8C), one can immediately see that to achieve this goal, the patient should be given the highest possible non-toxic dose. Since under some circumstances, the disease burden suggests a low-intermediate dose (and intermediate treatment coverage), there can hence be a conflict between the interest of the individual patient and of the population as a whole.

In the present article, the disease is non-lethal for all patients, even in the absence of treatment. For diseases with a potentially fatal outcome, the reduction in mortality would be a highly relevant population-level criterion. At the level of the individual, the within-host probability of resistance would be important since the evolution of resistance would not simply delay recovery but substantially increase the risk of death. A risk of pathogen-induced mortality could be included into our model, e.g. by introducing a risk of lethality beyond a certain pathogen load. The dose-choice strategies would then have to account for the necessity of containing the pathogen load below potentially-lethal levels [40].

We assume that all patients are immuno-competent. However, some hosts can be immunocompromised and therefore be particularly vulnerable to treatment failure. Even if the disease is non-lethal to the majority of the population, it could pose a major risk to these patients who are hence extremely dependent on the availability of efficient drugs. To protect them, suppressing the spread of resistance in the population would be a priority, and population-level measures of resistance emergence and spread would be very relevant. The drug dose administered to the immuno-competent part of the population should then optimally limit the spread of resistance even at the cost of delayed recovery. Obviously, higher doses would be required to treat the immuno-compromised patients.

### The concept of the therapeutic window

Whenever the relationship between the criterion used for the dose assessment and the drug dose itself is non-monotonic, whether the criterion shows a single peak (e.g. *p_e_*) or several local extrema (disease burden), any recommendation on the best dose is strongly dependent on where the therapeutic window lies. The therapeutic window depends on the minimum effect of treatment and the maximum of side effects that is considered acceptable. In our model, the recovery rate changes substantially across the therapeutic window. Besides clearing the pathogen, treatment could in addition directly mitigate symptoms. This would be another component of treatment effect. If low doses are similarly good at mitigating symptoms as high doses, the effects of treatment could be considered more similar at low and high doses despite differential recovery rates (however, in our model, symptoms are directly linked to the pathogen load).

It is quite likely that the therapeutic window varies between patients. Adjusting prescribed doses to the body weight of each patient is a way to partially account for host heterogeneity. Still, acquiring accurate knowledge on the therapeutic window for each patient may be challenging. Prescribing low-dose treatments can be dangerous if the lower bound of the therapeutic window is underestimated for a given patient. Here, the disease modeled is self-limiting and all hosts are immuno-competent such that under-dosing would not be life-threatening, but it will be for many diseases. The same can be true for overdosing.

What is relevant for our study (as it is for others) is the relationship between drug dosage and the in-vivo probability of resistance evolution within the therapeutic window. It is hard to obtain this information for a given disease and drug. Yet, clarifying this relationship can help to guide the development of new treatment strategies.

### Optimal treatment strength at the population level: Similarities between drug dosage, coverage, and population-wide timing of treatmen

The strength of treatment can be modulated in various ways. One of them is the choice of the drug dose as considered in the present study. At the population level, it is also possible to vary the fraction of the population that receives treatment (“coverage” or “treatment level”) and the optimal time point during the course of the epidemic when antimicrobials start being used in the population. The optimal choice of these two factors, given the risk of resistance evolution, has been assessed in a series of studies on optimal drug use during influenza epidemics (e.g. [33–35,38,39,41]).

There are many similarities between the findings of these studies and the present article. First, high treatment coverage right at the start of the epidemic might contain the spread of the disease, and so does high dose treatment in our model. More subtle is the case, when containment is impossible or has failed. If the number of susceptible hosts is a limiting factor to the disease spread - as it is in our simulated study and as it can be in real-life epidemics (e.g. influenza) -, the disease dynamics in the population is strongly affected by competition between the sensitive and the resistant strains for susceptible hosts. By choosing the right treatment strength (which may refer to drug dose, coverage, or timing of treatment), this competition may be exploited to reduce the overall negative impact of the disease on the population. Essentially, treatment needs to be strong enough to impeach the wide spread of the sensitive strain but weak enough to allow enough individuals to receive immunity through infection with the sensitive strain; then, spread of the resistant strain is hampered due to the reduced pool of susceptibles. If the strength of treatment is too strong, so-called “overshooting” occurs. Too many susceptibles remain untouched by the sensitive strain to establish herd immunity that would prevent or at least mitigate a time-delayed resistant epidemic [42]. The resistant strain then profits from competitive release.

With treatment strength referring to drug dose, this competitive release is translated in an optimal spread of resistance at high-intermediate doses that efficiently suppress the sensitive but not the resistant strain (cf. the prediction in [5]). The disease burden and the total number of infecteds therefore display a maximum in this dose range. In some low-intermediate dose range, where the sensitive strain is suppressed well but not well enough to allow for competitive release, we find a local minimum. Likewise, intermediate coverage or a delay in the use of antimicrobials in the population may be best in reducing the total number of cases [33–36,38,39]; see also our Fig 6. However, if the resistant strain suffers a strong transmission cost (i.e. *β* is much lower for the resistant strain), high coverage yields the best outcome [33,35]. We have not implemented such a transmission cost into our model, which might alter results. Another factor that has been considered in the context of influenza epidemics is prophylactic treatment, which – once the resistant strain has appeared in the population – substantially facilitates its spread since it meets little or no competition for the treated hosts with the sensitive strain [33, 41].

### Limitations and extension

The interactions between pathogen load, treatment, immune response, and symptoms are complex and depend on the disease and the drug. In our model, symptoms set in once the pathogen load reaches a given threshold. This implies that they are directly caused by the pathogen. However, symptoms can also be due to the action of the immune system, e.g. fever can be part of the immune response. Within our setup, this would mean that the onset of therapy would depend on the level of the immune response, *I*, rather than on the pathogen load, *P_w_* + *P_m_*. In that case, the drug dose would have less effect on the strength of the immune response. On the other hand, drugs can also have a direct effect on the immune system, which would be dose-dependent. For example, they can mitigate corresponding reactions such as fever. Both factors would change the relationship between dose and within-host probability of resistance evolution and hence conclusions on optimal dosing. On the other hand, resource competition also leads to an inverted U-shaped relationship between the within-host probability of resistance and the drug dose [12]. It is hence possible that the inverted U-shape persists even with different interactions between treatment and the immune system, if strain competition is taken into account, such that our results would carry over.

We described the immune response by a single variable *I* and do not separately model the innate and the adaptive immune system. We assumed that upon recovery, each patient has built up an immunological memory that protects the individual from reinfection at least until the end of the epidemic. This happens independently of the level of immune response, *I*, reached during the infection. However, with a high drug dose that quickly suppresses the pathogen, no strong immune response might be triggered and no (or only weak) immunological memory might develop. In that case, immunity (which would then be weak anyway) is fading during the time span of the epidemic, and reinfection cannot be excluded. The pathogen would then potentially be maintained in the population for a longer period of time, and the disease burden could be higher for aggressive therapy than predicted by our model. To investigate this, a more accurate modeling of the immune system would be necessary, following e.g. Ankomah and Levin (2014) or Gjini and Brito (2016) [28,29].

Our within-host model also ignores classical ecological factors such as spatial structure and temporal changes in drug pressure. Drug penetration is not equal across body tissues, potentially creating compartments with high and low drug concentrations. This spatial drug heterogeneity has major implications for the evolution of drug resistance [43–45], and it will also affect the dose-dependent recovery rate. Likewise, changes in the drug concentration upon drug intake and subsequent decay (pharmakokinetics) might have an effect on the within-host dynamics of the two strains. In this context, it would also be interesting to include incomplete adherence to treatment, where patients skip doses or stop treatment prematurely. Omitted doses could lead to a lower “effective dose”, i.e. aggressive chemotherapy might, for example, resemble treatment with intermediate doses.

Another set of assumptions concerns the between-host level. The mode of pathogen transmission assumes transmission of one strain only during a transmission event, even when the infecting host carries both pathogen strains. This assumption is justified if the two strains are not well-mixed inside the disease-carrying host and if the mode of transmission is likely to produce an inoculum consisting of pathogens that were originally grouped together. Alternatively, in other cases, transmission of a mixed inoculum containing pathogens drawn randomly in proportion with the load of each strain in the infecting host may be more realistic. This would probably alter the speed of spread of resistance in the population and mitigate strain competition, presumably leading to different results regarding optimal drug-dose choices. Another important assumption with consequences for the spread of resistance is the assumption of full cross-immunity between strains.

In our model, the transmission coefficient remains constant throughout the course of the infection. However, in many cases, patients will stay at home or be isolated, once symptoms set in. Another potential extension of our work would therefore be to include a diminished transmission coefficient for individuals with declared symptoms.

We considered a homogeneous and well-mixed population. Variation in the number of contacts (e.g. “superspreaders”) and more generally the contact structure of the social network influences the disease dynamics (see for instance [46,47]). This structure moreover affects the invasion of a second – for us, the resistant – strain [48]. It would be interesting to investigate how different contact structures affect optimal drug dosage. It might also turn out to be optimal to use different doses for different people, depending on their position in the network and their number of contacts. Individuals moreover differ in their immune response, as mentioned above.

For the antibiotic treatment of bacterial infections, bystander selection of resistance in the commensal flora is an important aspect and may be dose dependent [49–51]. Assessing the relationship between dose and resistance evolution in the commensals is beyond the scope of our study but of high practical relevance.

## Conclusion

Choosing the optimal drug dose within the therapeutic window can prove non-trivial, and the right decision depends on the measure of treatment success. Epidemiological parameters that are dose dependent (such as the period of infectiousness) influence the disease dynamics at the population level, bringing the problem of optimal dosage from the individual to the population. Importantly, both the within-host and the between-host dynamics affect the evolutionary dynamics of the pathogen and hence selection for resistance. For the management of the disease, both scales need to be taken into account.

## Method

### Model descriptio

We build a nested model of within-host and between-host dynamics. It is constructed by adopting the two-strain within-host model for acute, self-limiting infections from Day and Read, 2016 [12], and combining it with simple transmission rules for the between-host interactions (see Fig 7A for a typical course of infection and Fig 7B for a schematic representation of the nested model).

**Fig 7.**
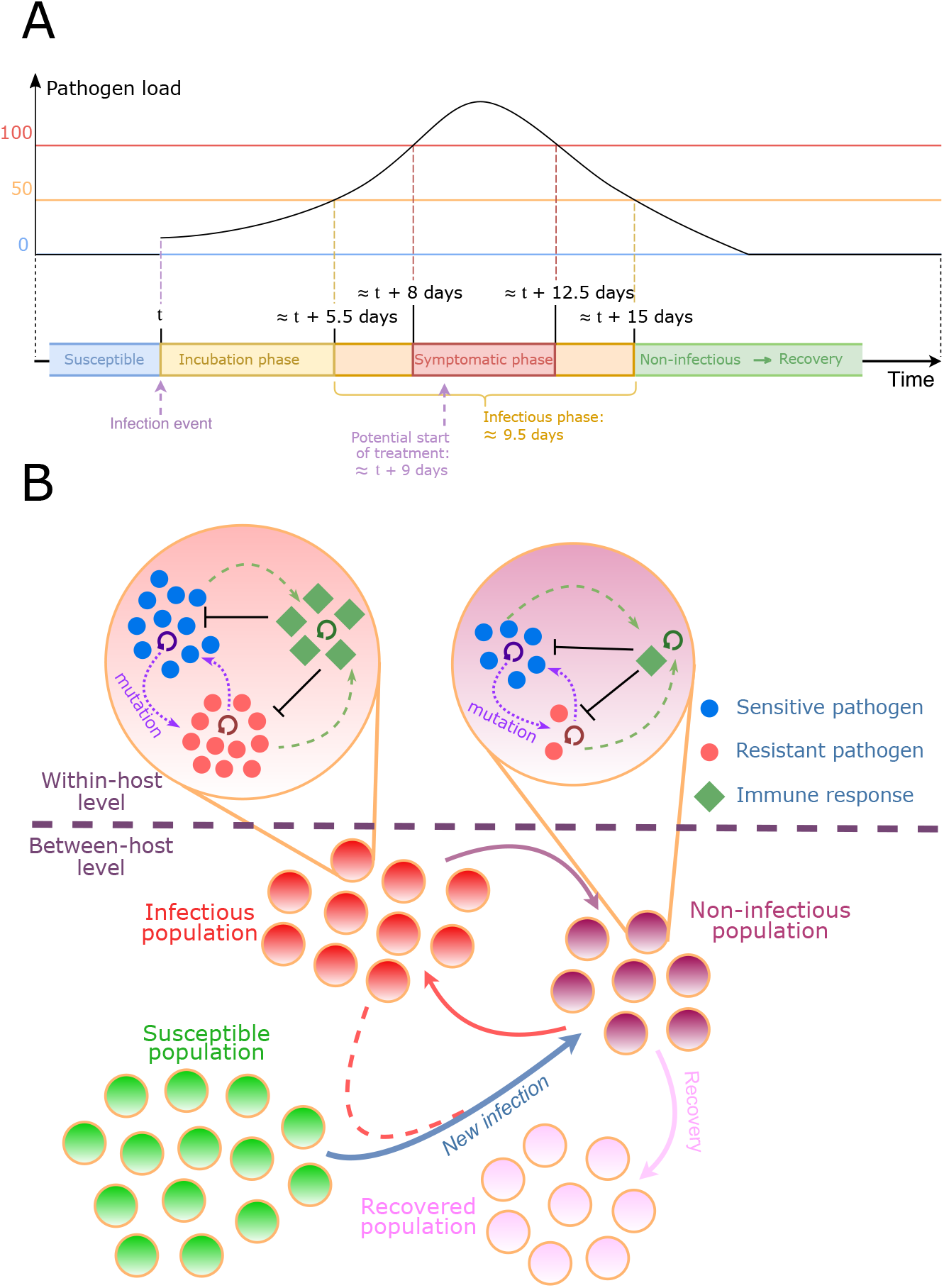
Nested model of an acute infectious disease. (A) Average timeline of the infection for a non-treated individual. The host’s transition from one phase to the other is determined by the pathogen load. (B) Diagram of the nested model.

### Within-host dynamic

The infecting pathogen can be found in the infected host under two different forms: a wild-type strain, and a mutant strain that withstands considerably higher drug pressure. Even in the absence of treatment, the host’s immune system is able to fully clear the infection. A treatment can be prescribed to reduce the period of infection.

The load of each pathogenic strain inside the host is denoted by *P_w_* and *P_m_* for the wild-type sensitive strain and the mutant resistant strain, respectively. Mutations happen in both directions during pathogen replication with probability *μ*. The drug acts by reducing the rep
lication rate of the pathogen, and the replication rates of each strain, *r_w_*(*c*) and *r_m_*(*c*), are decreasing functions of the drug concentration *c*. We assume that the drug concentration is constant throughout the course of treatment, neglecting any pharmacokinetics. The resistance mutation entails a cost, manifested in a lower replication rate in the absence of the drug (*r_w_* (0) ≥ *r_m_* (0)). Pathogens of both genotypes have an intrinsic death rate *η*.

The immune response, *I*, is triggered proportionally to the total pathogen load (*P_w_* + *P_m_*) with a factor λ and decays at a constant rate *δ*. The depletion of pathogens due to the immune system follows mass-action kinetics and leads to additional pathogen death rates *κP_w_I* and *κP_m_I*. Both strains are hence equally affected by the immune system, and there is full cross-immunity. As mentioned above, we choose the parameters of the model such that the disease is self-limiting, i.e., even in the absence of treatment, the infection is successfully suppressed by the host’s immune system.

Deterministically, the dynamics can be described by the following set of differential equations:

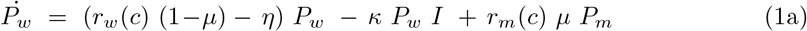

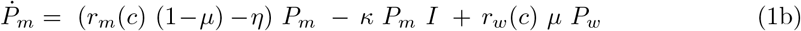

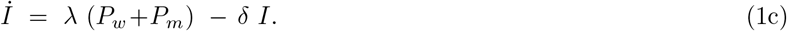

The explicit formulas for the replication rates *r_w_* and *r_m_* are given by:

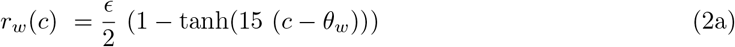

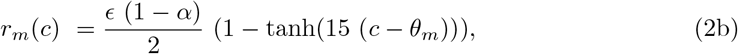

where *є* is the growth rate of the susceptible strain in the absence of treatment, *α* represents the fitness cost of resistance, *θ_w_* (*θ_m_*) is the dose for which the growth rate of the sensitive (resistant) strain is half the growth rate in the absence of treatment. Resistance is characterized by *θ_m_* > *θ_w_* and *α* ≥ 0.

Fig 7A shows the timeline of an average infection. A susceptible host gets initially infected by a small number of pathogens (at the same time, we choose this number high enough to make stochastic extinction unlikely; concretely, it is set to 7 pathogens). The pathogen load then increases during a symptom-free incubation period. Once it crosses a threshold of 50 pathogens, the infected individual turns infectious, and once the pathogen load crosses a threshold of 100 pathogens, symptoms set in. Treatment is applied after a delay of one day, which subsumes all steps between the appearance of symptoms and the moment when treatment starts being effective. The drug is then administered for a fixed period of 7 days and renewed as many times as necessary to suppress the pathogen load below the symptoms threshold. In the unlikely case that the pathogen load increases again to cause symptoms, therapy is resumed. If the pathogen load of an infected individual has dropped to zero, this individual is considered as recovered and is assumed to have acquired life-long immunity to the disease due to immunological memory, which we do not model explicitly. We choose the parameters such that in a typical course of infection, patients become infectious about 5.5 days after infection, and develop symptoms 2.5 days later. We choose *μ* = 0.001, *η* = 0.02 day^-1^, *κ* = 0.0175 day^-1^, *λ* = 0.035 day^-1^, *δ* = 0.01 day^-1^, *є* = 1.2 day^-1^, *α* = 0.017, *θ_w_* = 0.3, *θ_m_* = 0. 6.

### Between-host dynamic

Initially, a single individual is infected with the sensitive wildtype strain, and all other individuals are susceptible. Since the infection is assumed to be non-lethal and since the duration of the epidemic outbreak is short for the vast majority of parameter sets used in this study (in the order of 10^2^ days), we ignore birth and death of individuals as well as immigration into or emigration from the population. The total population size is set to *N* = 10,000 throughout the manuscript.

For simplicity, we assume that all infectious individuals are equally likely to infect another population member, independent of pathogen load, genetic composition of the pathogen population, and treatment status. There is therefore no direct transmission cost of the resistant strain and chances of transmission depend on the infecting host’s pathogen load in a step-function fashion (no transmission below a certain threshold, full transmission above). All individuals (infected or not) except recovered ones are potential targets for infection, and infection occurs randomly between individuals, i.e. we disregard population structure. With *I*_inf_ the number of infectious hosts, *R* the number of recovered individuals, and *β* the transmission coefficient, the rate at which infection events happen in the population is given by *β* × *I*_inf_ × (*N − R −* 1). The “−1” accounts for the fact that infectious individuals do not infect themselves.

The inoculum size at infection is fixed to 7 pathogens and again independent of the infecting host’s pathogen load. If the infecting host is co-infected by both pathogen strains, the transmitted strain is chosen randomly in proportion to the pathogen load of each strain in this host. Only one strain is therefore transmitted at each infection event. This relies on the idea that the two pathogen strains are not well mixed in a co-infected individual and that pathogens in close spatial proximity are likely to be transmitted together, which in turn makes it likely that all pathogens in the inoculum derive from the same clone. Superinfection of an already infected individual follows the same rules as infection of a pathogen-free individual.

Within this framework, the epidemic occurs in one “wave” and then goes extinct. Even for high transmission coefficients, persistence is impossible since, as the epidemic proceeds, the number of susceptible hosts drops to a level too low to maintain the epidemic. For low transmission coefficients, for which the average number of secondary cases is less than one, most outbreaks will be small. For high transmission coefficients that are large enough for the epidemic to spread, the distribution of outbreak sizes is bi-modal. Either the epidemic comes to a halt quickly due to stochastic extinction, or a large outbreak occurs that is only stopped by the lack of a sufficient number of susceptible hosts (see Fig S1).

## Analysi

### Agent-based simulation

We run stochastic simulations of the nested model of within-host and between-host dynamics. The differential equations of the within-host level (Eq 1) and the state-transition rules for the between-host level are implemented stochastically, using the fixed-step-size tau-leaping Gillespie algorithm [52,53]. Simulations start at time *t* = 0 with one individual newly infected with the wild-type strain of the pathogen. At each time step, the within-host infection dynamics of all the infected individuals are updated, using the stochastic counterpart of Eq 1 with exponentially distributed waiting times. At the end of each step, transmission events (new infections and superinfections) are randomly drawn, using the rules defined in the previous paragraph. Simulation runs end when there is no pathogen-carrying host in the population anymore. In all simulations presented here, the time steps of the algorithm are of length *δ*_t_ = 5 × 10^-3^ days. The implementation is done in Java.

### Simplification into a deterministic SIR mode

The agent-based nested model is complex to analyze and to dissect. To gain a better and more intuitive understanding, we introduce a classical epidemiological model, representing a simplified version of the dynamics. Except for the transmission coefficient *β*, all the parameters of this simplified model are extracted from simulations of the stochastic within-host dynamics of the nested model. The simplified model allows us in particular to efficiently investigate the influence of the coefficient of transmission of the epidemic *β*, which is shared between the two models.

For the simplified deterministic model, we divide patients into four groups. Individuals can be either susceptible, infected by one of the two strains, or recovered (SIR model). As we will see below, the clear classification of infections as sensitive or resistant provides particularly helpful insight into the interaction of both strains at the between-host level. The relevant processes, leading to transitions between the compartments, are infection, patient recovery, and the emergence of resistance. We describe the dynamics by a set of deterministic differential equations:

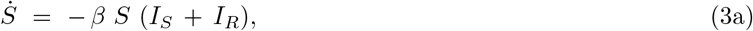

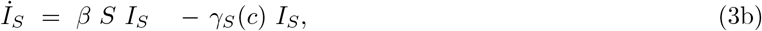

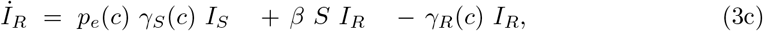

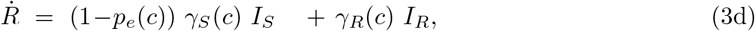

where *S* is the compartment of susceptible hosts, *I_S_* and *I_R_* the compartments of infected hosts, respectively with the sensitive and resistant strain, and *R* is the compartment of recovered individuals. As before, *β* is the transmission coefficient of the pathogen, assumed identical for both strains. *γ_S_*(*c*) and *γ_R_*(*c*) are the dose-dependent recovery rates of individuals infected with the sensitive strain and the resistant strain, and *p_e_*(*c*) is the probability of resistance emergence at the within-host level. We estimate the functions *γ_S_*(*c*), *γ_R_*(*c*), and *p_e_*(*c*) from simulations of the within-host model introduced above (Fig 8). Details are given in SI section S3.1.

**Fig 8.**
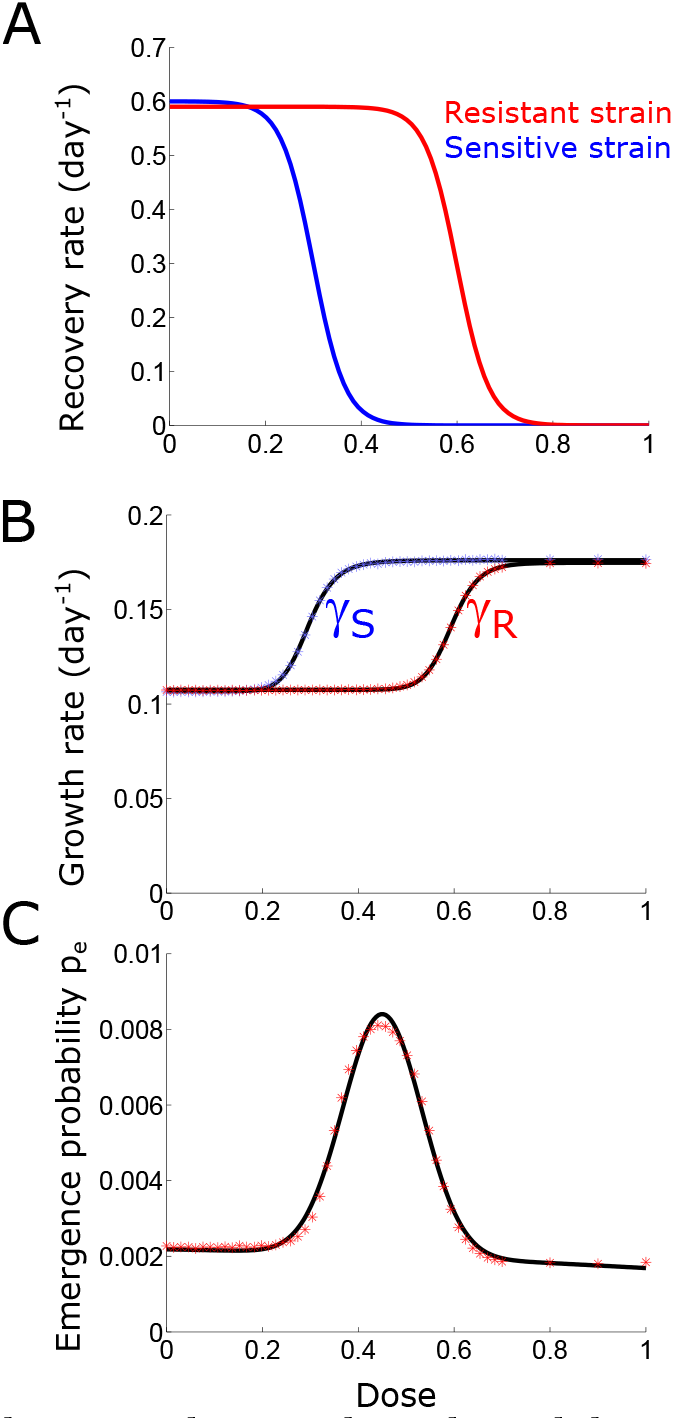
Transition from the nested agent-based model to an SIR model. (A) Effect of treatment strength on the growth rate for the sensitive and resistant strains. At very high doses, neither strain can grow in the presence of treatment. Parameters: *є* = 0.6, *α* = 0.017, *θ_w_* = 0.3, *θ_m_* = 0.6. (B) Mean recovery rates of an individual infected by the sensitive (blue) and resistant (red) strain. Simulated results are fitted with sigmoid functions. (C) Probability of emergence of resistance as a function of dose, obtained from 10 million within-host simulations per dose (red) and fitted with the sum of two Gaussians (black).

The SIR model provides a deterministic non-nested approximation of the full agent-based model. It only describes the between-host dynamics and therefore does not allow for feedbacks between the within-host and between-host levels. While differing in many respects from the agent-based model, the SIR model is still able to capture important aspects of the dynamics such as competition of the two pathogen strains for susceptible hosts at the population level. While we focus on the deterministic SIR model in the main text, we also discuss the relation of the agent-based model to the stochastic SIR model (i.e. the stochastic version of Eq 3) in SI section S3.4.

For the SIR model, the basic reproductive number (i.e. the number of secondary infections caused by one infected individual in a fully susceptible population) of the sensitive strain in the absence of treatment can be obtained as *R*_0_ = *βN*/*γ_S_*(0). As *R*_0_ is a very informative quantity in epidemiology - in particular, an epidemic can spread with non-zero probability for *R*_0_ > 1 and quickly goes extinct otherwise -, we will in the following report this *R*_0_, when we vary *β*. In the analysis presented here, the “true” basic reproductive number in the agent-based model hardly differs from the basic reproductive numbers calculated for the SIR model (Fig S4).

### Evaluating the success of a treatment strateg

Various criteria can be used to assess the quality of a treatment strategy, which - in our context - refers to the choice of drug dosage. We use here four different measures. The first two consider overall treatment success, while the other two focus on the evolution and spread of resistance.

The first two criteria are based on the disease burden, which we define as the cumulative number of days that members of the population spend being infectious. With *δ_t_* the step-size of the tau-leaping algorithm, and *I*(*t_i_*) the number of infectious hosts in the *i^th^* calculation step, and *K* the number of steps of the algorithm needed to reach the disease-free equilibrium, *B*_0_ is approximated as:

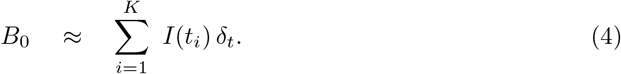

We focus on infectious individuals rather than individuals showing symptoms to allow for a comparison of results from the agent-based model and the SIR model. As discussed below, this does not alter the conclusions.

### Probability of an epidemic outbrea

Initially, only a single individual is infected. It is therefore possible that due to stochasticity, the epidemic dies out after only a few infection events rather than developing into a serious outbreak with many infecteds. We use the probability of an epidemic outbreak as a first measure of success of a chosen drug dose. To evaluate concretely if an epidemic has broken out in a simulation run, we focus on the disease burden *B*_0_.

As pointed out above and shown in Fig S1, the distribution of *B*_0_ is bimodal unless the transmission coefficient is low and the drug dose high. In the vast majority of cases, a threshold of 300 patient days clearly separates epidemics that quickly went extinct due to stochasticity from serious outbreaks. Thus, we define the outbreak probability as the probability that the disease burden reaches at least 300 patient days and determine it as the fraction of simulation runs which yield a disease burden larger than the 300-patient-days threshold.

As is intuitive and confirmed by simulations (not shown), the outbreak probability is independent of whether we focus on infectious or symptomatic cases.

### Total disease burden over the course of an epidemi

As a second criterion, we assess the severity and magnitude of spread of an epidemic measured by the disease burden, provided an outbreak occurs, i.e. we only consider epidemics where the disease burden reaches at least 300 patient days. We denote the average disease burden in this subset of epidemics by *B*. This provides a measure of the overall negative impact that an epidemic has on the population. In the following, when we speak about the disease burden, we mean *B* (rather than *B_0_*).

In the SIR framework, the disease burden is given by

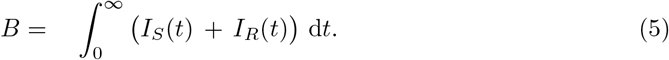

Note that since the SIR model is deterministic, the disease certainly spreads when one of the strains has a basic reproductive number > 1, while the number of the infecteds cannot increase at all otherwise. We therefore do not include any conditioning.

### Probability of emergence of resistance at the within-host leve

Focusing on the evolution of resistance, one aim is to minimize the probability of *de novo* emergence of resistance within a host, *p_e_*, as is done by Day and Read (2016) [12]. We say that resistance has emerged if the pathogen load crosses a given threshold. As was done by Day and Read with quite similar parameters for the within-host model, we set this threshold to 100 resistant pathogens, which corresponds to the threshold for the appearance of symptoms.

### Spread of resistance at the between-host leve

As a measure of the risk of emergence and spread of resistance in the population, we count the number of transmission events of the resistant strain from one host to another (remember that only a single strain is transmitted at infection). These transmission events can either be superinfections of already-infected individuals or infections of susceptible hosts. Restricting this measure to infection of susceptible-hosts-only yields qualitatively similar results.

## Acknowledgements

The authors thank Sebastian Bonhoeffer, Roland Regös, Joachim Hermisson for helpful discussions. The authors are grateful to Pleuni Pennings and Arne Traulsen for valuable comments on the manuscript.

## Supporting information caption

**S0 File**. Full supporting information (text and figures)

